# Prediction performance of linear models and gradient boosting machine on complex phenotypes in outbred mice

**DOI:** 10.1101/2021.08.02.454826

**Authors:** B.C. Perez, M.C.A.M. Bink, G.A. Churchill, K.L. Svenson, M.P.L. Calus

## Abstract

Recent literature suggests machine learning methods can capture interactions between loci and therefore could outperform linear models when predicting traits with relevant epistatic effects. However, investigating this empirically requires data with high mapping resolution and phenotypes for traits with known non-additive gene action. The objective of the present study was to compare the performance of linear (GBLUP, BayesB and elastic net [ENET]) methods to a non-parametric tree-based ensemble (gradient boosting machine – GBM) method for genomic prediction of complex traits in mice. The dataset used contained phenotypic and genotypic information for 835 animals from 6 non-overlapping generations. Traits analyzed were bone mineral density (BMD), body weight at 10, 15 and 20 weeks (BW10, BW15 and BW20), fat percentage (FAT%), circulating cholesterol (CHOL), glucose (GLUC), insulin (INS) and triglycerides (TGL), and urine creatinine (UCRT). After quality control, the genotype dataset contained 50,112 SNP markers. Animals from older generations were considered as a reference subset, while animals in the latest generation as candidates for the validation subset. We also evaluated the impact of different levels of connectedness between reference and validation sets. Model performance was measured as the Pearson’s correlation coefficient and mean squared error (MSE) between adjusted phenotypes and the model’s prediction for animals in the validation subset. Outcomes were also compared across models by checking the overlapping top markers and animals. Linear models outperformed GBM for seven out of ten traits. For these models, accuracy was proportional to the trait’s heritability. For traits BMD, CHOL and GLU, the GBM model showed better prediction accuracy and lower MSE. Interestingly, for these three traits there is evidence in literature of a relevant portion of phenotypic variance being explained by epistatic effects. We noticed that for lower connectedness, i.e., imposing a gap of one to two generations between reference and validation populations, the superior performance of GBM was only maintained for GLU. Using a subset of top markers selected from a GBM model helped for some of the traits to improve accuracy of prediction when these were fitted into linear and GBM models. The GBM model showed consistently fewer markers and animals in common among the top ranked than linear models. Our results indicate that GBM is more strongly affected by data size and decreased connectedness between reference and validation sets than the linear models. Nevertheless, our results indicate that GBM is a competitive method to predict complex traits in an outbred mice population, especially for traits with assumed epistatic effects.

## INTRODUCTION

The use of genome-wide markers as predictor variables for individuals’ unobserved phenotypes (Meuwissen et al., 2001) based on a reference population is known as genomic prediction (GP). In the past decade, high-throughput genotyping technologies made GP accessible and facilitated large-scale use of GP for animal (Boichard, 2016) and plant (Bhat et al., 2016) breeding, and in human genetics (Lappalainen et al., 2019). For animals and plants, GP has reduced breeding costs and speeded up breeding programs as individuals of interest can be selected in earlier stages of life, while reducing costs for performance testing. In humans, major efforts have been put into developing GP to score disease risks (Duncan et al., 2019), aiming for a more personalized medicine in the future (Barrera-Saldaña, 2020).

Currently, most GP models implemented assume that observed phenotypes are controlled by numerous loci with additive effects throughout the genome and this approach has provided a robust performance in most cases (Meuwissen et al., 2001; Calus, 2010). However, in the literature it has been suggested that the genetic architecture of complex traits may involve significant proportions of non-additive genetic (dominance or epistasis) effects (Mackay, 2014) and that these could be much more common than previously thought (Sackton and Hartl, 2016). Although accounting for non-additive effects into parametric GP models has been reported to improve predictive performance (Forsberg et al., 2017) of phenotypes, implementing variable selection to prioritize among all possible SNP by SNP interactions, is computationally too costly for any practical application.

Machine learning (ML) has been successfully used in many fields for text, image and audio processing at huge data volumes. Recently, these algorithms have found many applications in GP for offering an opportunity to model complex trait architectures in a much simpler framework than parametric models (Nayeri et al. 2019; Montesinos-López et al., 2021; van Dijk et al., 2021). ML algorithms are free from model specification, can accommodate interactions between predictive variables and deal with large numbers of predictor variables by performing automatic variable selection (Jiang et al., 2009; Li et al., 2018).

Howard et al. (2014), Ghafouri-Kesbi et al. (2015) and Abdolahi-Arpanahi et al. (2020) have compared the predictive performance of linear and ML models for simulated phenotypes controlled by additive or non-additive effects. In general, linear models were able to outperform ML models for traits controlled by additive effects, however they failed to do so when used to predict traits with purely epistatic architecture. The superiority of ML over traditional linear models was markedly observed for traits controlled by a low number of loci (100) with non-additive effects. For this type of scenario, Ghafouri-Kesbi et al. (2015) and Abdolahi-Arpanahi et al. (2020) also showed a consistent good performance of the gradient boosting machine (GBM) algorithm (Friedman, 2001), which has previously been reported to provide robust predictive ability when compared to other methods in the context of GP (González-Recio et al., 2011, 2013, 2014; Ogutu et al., 2011; Jimenez-Montero et al., 2013; Grinberg et al., 2019; Srivastava et al., 2021).

Although results in simulated data suggest the superiority of ML models in the presence of epistatic effects, the performance of such models have been much less consistent for GP using real datasets. Zingaretti et al. (2020) observed that convolutional neural networks (CNN) had 20% higher predictive accuracy than linear models for GP of a trait with a strong dominance component (percentage of culled fruit) in strawberry but underperformed for traits with predominant additive effects. On the other hand, in Azodi et al. (2019), ML did not consistently outperform linear models for traits with strong evidence of underlying non-additive architectures (for example height in maize and rice). The authors also describe that ML models presented less stable prediction across traits than linear models. Similar results were also reported by Bellot et al. (2018) while investigating the performance of GP for several complex human phenotypes. An important aspect to consider when investigating performance of GP models is that for most livestock and plant species there is currently limited knowledge over the genetic architecture of economically interesting traits. This makes it difficult to perform inference about the real reasons why ML outperforms linear models in specific situations. This could be overcome by considering data from populations for which knowledge on genetic architecture of traits is more extensively and accurately described.

The Diversity Outbred (DO) mice population is derived from eight inbred founder strains (Svenson et al. 2012). It is an interesting resource for high-resolution genetic mapping by having a low level of genetic relationship between individuals, low extent of LD (Churchill et al., 2012) and uniformly distributed variation across genomic regions of known genes (Yang et al., 2011). This structure represents an advantage over classical inbred strains of mice or livestock populations, which have limited genetic diversity (Yang et al. 2011). These aspects allow the investigation of relevant traits in a structured scheme that closely reflects the genetic mechanisms of human disease (Churchill et al., 2012, Svenson et al., 2012).

In the present study, the objective was to compare performance of GBM to several linear models (GBLUP, BayesB and elastic net) for predicting ten complex phenotypes in the DO mice population. All models were applied for scenarios where data was not available for one or more generations in between the reference and validation sets. Additionally, we explore the use of feature selection from the GBM algorithm as a tool for sub-setting relevant markers and to improve prediction accuracy through dimensional reduction.

## MATERIAL AND METHODS

### Data

#### Phenotypes

The DO mice dataset comprising 835 animals was obtained from The Jackson Laboratory (Bar Harbor, ME). The animals originated from 6 non-overlapping generations (4, 5, 7, 8, 9 and 11) in which males and females were represented equally. The total number of animals per generation was 97, 48, 200, 184, 99 and 197 for generations 4, 5, 7, 8, 9, and 11, respectively, but numbers of missing records varied across traits (Figure 1). The mice were maintained on either standard high fiber (chow, n=446) or high fat diet (HFD; n=389) from weaning until 23 weeks of age. The proportion of males and females within each diet category was close to 50-50 for all generations. The same was observed for the frequency of males and females within each litter-generation combination (two litters per generation). A detailed description of husbandry and phenotyping methods can be found in Svenson et al. (2012).

**Figure 1.**
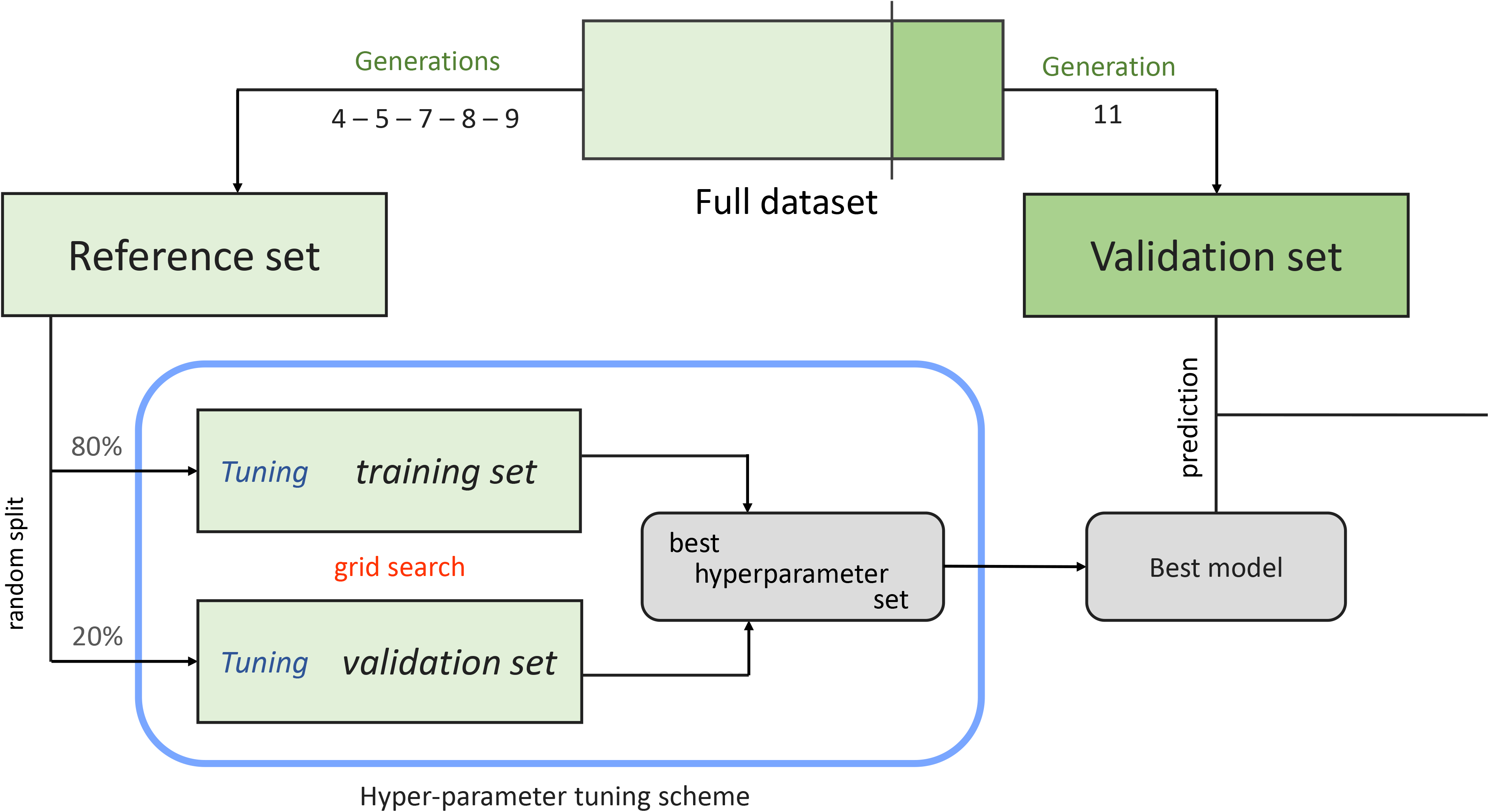
Graphical representation of the hyper-parameter tuning grid-search scheme implemented to obtain the best GBM and ENET models.

Table 1 shows a comprehensive description of each trait regarding dataset size, estimated heritability and assumed genetic architecture with associated literature. Among all phenotypes available we chose 10 traits based on their distinct assumed genetic architectures from previous results with the same dataset (Li and Churchill, 2010; Churchill et al.,2012; Zhang et al., 2012; Tyler et al.,2016, 2017; Keller et al., 2019; Keenan et al., 2021) and other populations (Chitre et al., 2018). The analyzed traits were bone mineral density at 12 weeks (BMD), body weight at 10, 15 and 20 weeks (BW10, BW15 and BW20); circulating cholesterol at 19 weeks (CHOL), adjusted body fat percentage at 12 weeks (FATP), circulating glucose at 19 weeks (GLU), circulating triglycerides at 19 weeks (TRGL), circulating insulin at 8 weeks (INSUL) and urine creatinine at 20 weeks (UCRT). These traits can be categorized into measurements of body composition (weights and fat percentage), clinical plasma chemistries (triglycerides, glucose, insulin) and urine chemistry (urine creatinine).

**Table 1.** Number of available observations (N), estimated heritability, assumptions from literature regarding the genetic architecture of the trait and references.

| Trait | N | Heritability | Genetic Architecture | Reference |
| --- | --- | --- | --- | --- |
| BMD | 831 | 0.39 | Evidence of epistatic effects | Tyller *et al.* (2016) |
| BW10 | 834 | 0.42 | Highly polygenic | Tyller *et al.* (2017)<br>Chitre *et al.* (2018) |
| BW15 | 829 | 0.34 | Highly polygenic | Tyller *et al.* (2017)<br>Chitre *et al.* (2018) |
| BW20 | 827 | 0.37 | Highly polygenic | Tyller *et al.* (2017)<br>Chitre *et al.* (2018) |
| FATP | 831 | 0.44 | Highly polygenic | Tyller *et al.* (2017) |
| CHOL | 819 | 0.33 | QTL with high effect<br>Evidence of epistatic effects | Stewart *et al.*, (2010)<br>Li and Churchill (2010)<br>Zhang *et al.* (2012) |
| GLUC | 816 | 0.12 | Evidence of epistatic effects | Stewart *et al.* (2010)<br>Chen *et al.* (2017) |
| INSUL | 820 | 0.21 | QTL with high effect | Keller *et al.* (2019) |
| TRGL | 820 | 0.29 | Highly polygenic | Stewart *et al.* (2010) |
| UCRT | 799 | 0.13 | Highly polygenic<br>Evidence of dominance effects | Perry (2019) |

<sup>1</sup> Standard error was close to 0.08 for all traits.

**Table 2.** Prediction error (mean squared error) obtained from GBLUP, BayesB, ENET and GBM for 10 phenotypes analyzed in the diversity outbred mouse population. Per trait, the lowest values are indicated in bold.

| Trait<sup>1</sup> | GBLUP | BayesB | ENET | GBM |
| --- | --- | --- | --- | --- |
| BMD | 0.886 | 0.929 | 0.904 | **0.885** |
| BW10 | **0.023** | 0.029 | 0.025 | 0.024 |
| BW15 | **0.025** | 0.030 | 0.026 | 0.025 |
| BW20 | **0.029** | 0.033 | 0.030 | 0.030 |
| CHOL | 0.068 | 0.104 | 0.080 | **0.066** |
| FATP | **0.486** | 0.523 | 0.488 | 0.493 |
| GLUC | 0.054 | 0.061 | 0.056 | **0.051** |
| TRGL | **1.339** | 1.503 | 1.373 | 1.367 |
| INSUL | **0.198** | 0.261 | 0.233 | 0.202 |
| UCRT | **0.019** | 0.022 | 0.020 | 0.020 |

<sup>1</sup>Bone mineral density at 12 weeks (BMD), Body weight at 10, 15 and 20 weeks (BW10, BW15 and BW20); circulating cholesterol at 19 weeks (CHOL), adjusted body fat percentage at 12 weeks (FATP), circulating glucose at 19 weeks (GLU), circulating triglycerides at 19 weeks (TRGL), circulating insulin at 8 weeks (INSUL) and urine creatinine at 20 weeks (UCRT)

Prior to any analyses performed in this study, phenotypic records were pre-corrected for fixed effects of diet, generation, litter and sex. The pre-corrected phenotype (*y*\*) can be represented by:

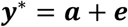

where ***a*** is the vector of animal additive genetic effects and ***e*** the vector of residuals.

#### Genotypes

Mice from 8 distinct founder strains were genotyped using either the MUGA and MegaMUGA SNP arrays (Morgan et al. 2016). The variant calls from the arrays in the animals contained in the current dataset were converted to founder haplotypes using a hidden Markov model (HMM) (Gatti et al. 2014), which uses the order of SNPs in an individual mouse to infer transition points between different DO founder haplotypes. After that, the probability of each parental haplotype at each SNP position in the genome (Gatti et al., 2014) was used to derive SNP genotype probabilities. To accomplish that, we used functions available in the “QTL2” R package (Broman et al. 2018). The complete genotype file used for the analyses was composed of 64,000 markers reconstructed from the diplotype probabilities from the MUGA and MegaMUGA on an evenly spaced grid, and the average distance between markers was 0.0238 cM. The full genotype data (64K markers) was cleaned based on the following criteria: variants with minor allele frequency < 0.05, call rates < 0.90 and linear correlation between subsequent SNPs > 0.98 were removed. After quality control, a total of 52,840 SNP markers were available for the mice with both phenotypic and genotypic records.

### Genomic prediction models

#### GBLUP

The statistical model of GBLUP is:

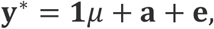

where **y**\* is the vector of pre-corrected phenotypes, **1** is a vector of ones, *μ* is the intercept, **a** is the vector of random additive genetic values, where 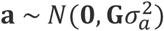 and **G** is the additive genomic relationship matrix between genotyped individuals. It is constructed following the second method described by VanRaden (2008) as 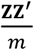 where **Z** is the matrix of centered and standardized genotypes for all individuals and *m* is the number of markers, and 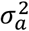 is the additive genomic variance, **e** is the vector of random residual effects where 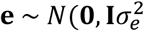 with 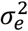 being the residual variance, and *I* is an identity matrix. GBLUP was implemented using a Bayesian approach using the BGLR package (Pérez and de los Campos, 2014). The Gibbs sampler was run for 150,000 iterations, with a 50,000 burn-in period and a thinning interval of 10 iterations. Consequently, inference was based on 10,000 posterior samples.

#### BayesB

BayesB has been widely used for genomic prediction (Meuwissen et al., 2001), and here we considered it for being a linear model with variable selection ability. The phenotype of the *i*^th^ individual is expressed as a linear regression on markers:

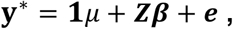

where **y**\* is the vector of pre-corrected phenotypes, **1** is a vector of ones, *μ* is the intercept, ***β*** is the vector of random effect of markers, ***Z*** is the incidence matrix for markers and ***e*** is a random residual where 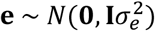 with 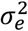 being the residual variance, and **I** is an identity matrix. Contrary to GBLUP, BayesB assumes *a priori* that all markers do not contribute to genetic variation equally. For BayesB, all markers are assumed to have a two-component mixture prior distribution. Any given marker has either a null effect with known prior probability, π, or a *t* prior distribution with probability (1 − π), with *v* degrees of freedom and scale parameter *S*^2^. Therefore, marker effects 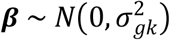, where 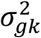 is the variance of the *k^th^* SNP effect. The BayesB model was implemented using the BGLR package (Pérez and de los Campos, 2014). The Gibbs sampler was run for 120,000 iterations, with a 20,000 burn-in period and a thinning interval of 100 iterations. Consequently, inference was performed based in 10,000 posterior samples.

#### Elastic Net

The elastic net (ENET) is an extension of the lasso (Friedman et al., 2010) and is considered a robust method under the presence of strong collinearity among predictors, as is the case for genotype data. It can be described by the regression model:

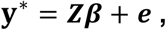

where **y**\* is the vector of pre-corrected phenotypes, ***β*** is the vector of random effect of markers, ***Z*** is the incidence matrix for markers and ***e*** is a random residual where 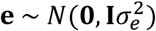 with 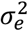 being the residual variance, and **I** is an identity matrix..

The ENET uses a mixture of the *ℓ*_1_ (lasso) and *ℓ*_2_ (ridge regression) penalties and the estimator 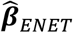 can be formulated as:

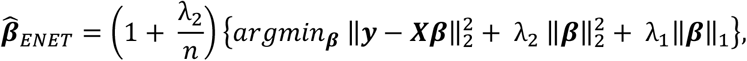

where 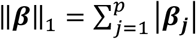 is the *ℓ*_1_-norm penalty on ***β***, 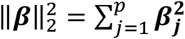 is the *ℓ*_2_-norm penalty on ***β***, 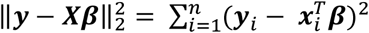 is the *ℓ*_2_-norm (quadratic) loss function (residual sum of squares), 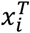 is the i-th row of ***X***, λ_1_ is the parameter that controls the extent of variable selection and λ_2_ is the parameter that regulates the strength of linear shrinkage. When setting 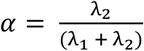, the ENET estimator is equivalent to the minimizer of: 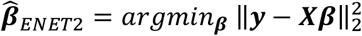, subject to 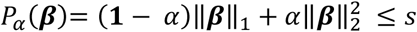 for some *s* where *P_α_* (***β***) is the ENET penalty (Zou and Hastie, 2005). The ENET is equivalent to ridge regression (Hoerl and Kennard, 1970) when *α* = 1, and to the lasso when *α* = 0. In practice, the *ℓ*_1_ component performs automatic variable selection while the *ℓ*_2_ component ensures that a group of highly correlated variables get effect estimates of similar magnitude.

We implemented the ENET model using the h2o.ai R package (Click et al. 2016). To establish the best hyperparameter set for ENET, we performed a cross-validation (splitting the reference set into 80-20 for train/test sets, as depicted in Figure 1) on a two-step scheme. First a grid search of values for the parameter *α* considering from 0 to 1, in intervals of 0.05. For tested value of *α*, the best value of λ was obtained by computing models sequentially, starting with λ = 1 and decreasing it exponentially until 0.01 in up to 20 steps. For each analysis, the best ENET model was chosen by the combination of *α* and λ parameters obtained from the grid search that yielded the lowest mean squared error of prediction in the test set, and this model was used to predict the validation animals (Supplementary Material - Figure S1).

#### Gradient Boosting Machine

Gradient boosting machine (GBM) is an ensemble learning technique that applies an iterative process of assembling “weak learners” into a stronger learner, being largely used for both classification and regression problems (Friedman, 2002;). It relies on fitting decision trees as the base learner (Hastie et al., 2009). The first tree is fitted on the errors of an initialized prediction based on the distribution of the response variable and from this point, the algorithm fits sequential trees, in which every subsequent tree aims to minimize the prediction error from the previous one until no further improvement can be achieved. Many different parameters can be used to measure that “improvement”, in the present study we used the mean squared error (MSE). GBM does automatic feature selection, prioritizing important variables and discarding ones containing irrelevant or redundant information. We implemented the GBM model using the h2o.ai R package (Click et al. 2016).

The performance of machine learning methods can be sensitive to hyper-parameters (Azodi et al., 2019). To obtain the best possible results from the GBM algorithm, a grid search approach was used to determine the combination of hyperparameters that maximized prediction performance for each trait. Hyperparameters (and range of values) included were number of trees (*ntree* = 100, 150, 200, 300, 500, 1000, 2000 and 5000), learning rate (*lrn_rate* = 0.01; 0.05 and 0.10) and maximum tree depth (*max_depth* = 2, 3, 5 and 10). For each trait analyzed, the hyperparameter tuning scheme was performed inside the reference subset (cf. ENET and Figure 1). The best set of hyperparameters was chosen based on the lowest mean squared error obtained from the grid-search. Results reported in the present study for GBM model refer to the best performing model out of the grid search for each trait (Supplementary Material - Figure S1).

#### Model performance

Performance of predictions from the models was measured by the accuracy, computed as the Pearson correlation 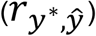, and the mean squared error of prediction (MSE) between predicted 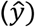 and pre-corrected phenotypes 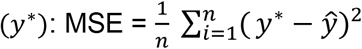 In all analyses, we used a forward prediction validation scheme in which animals from older generations (4, 5, 7, 8 and 9) were used as the reference and animals from the younger generation (11) as the validation subset. Uncertainties around the ***r** y*\*,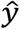 estimates were obtained by using bootstrapping (Davison and Hinkley, 1997), implemented in the “boot” R package (Canty and Ripley, 2021).

### Impact of the distance between a fixed-size reference and the validation set

Here we tested the impact of an increase in distance between the reference and validation sets on the prediction performance of each model. To accomplish that, we considered 3 scenarios using generation 11 as validation as before: Using generations 4, 5, 7, 8 and 9 as reference (NoGAP), using generations 4, 5, 7 and 8 as reference and omitting phenotypes from generation 9 (GAP9), using generation 4, 5 and 7 as reference and omitting phenotypes from generations 8 and 9 (GAP8+9). Considering the full dataset there were in total 638 animals from generations 4 to 9 available to be sampled for the validation subset. To analyze the proposed scenarios, the number of animals sampled for the reference subset was kept the same in all scenarios (N=300), with a constraint on the number of animals sampled from each generation to match its representativeness in NoGAP scenario (Supplementary Material - Table S2 for details). The fixed sample size of 300 was arbitrarily chosen based on the number of records available in GAP89, the scenario with the least available data to be sampled for the reference subset (N=345). Every scenario was evaluated in 20 replicates, inference was based on the average and standard deviation of accuracies obtained from replicates. All described models were applied to each of the 20 replicates (in every scenario) considering the same sampled dataset in each replicate across models. The complete list of animals sampled in each of the 20 replicates used for the analyses is provided in the Supplementary Material.

#### Feature importance for dimensionality reduction

For GBM, the importance of a feature is determined by assessing whether that feature was selected to split on during the tree building process, and the contribution of that to decrease the squared error (averaged over all trees) as a result (Friedman and Meulman, 2003; Hastie, Tibshirani and Friedman, 2009). The feature importance is expressed in a percentage scale that can be ranked to assess the magnitude of importance of each feature.

Here we investigate if the feature importance performed by the GBM model can be used to improve performance by fitting only extracted relevant features, i.e., SNPs, in GBM or any of the other models. We considered the top 100, 250, 500 and 1000 features from a GBM model using the cross-validation strategy previously explained as input for GBLUP, ENET and GBM models. The important features were obtained using the same strategy described for the hyperparameter tuning previously explained, thus using a random split (80-20) within the reference subset (Figure 1).

#### Similarities among top SNPs and prediction rankings

To assess the relationship between model’s prediction at the animal level, we quantified the number of animals in common in the top 20 ranked animals (approximately top 10% of generation 11) from each model. The latter metric gives an indication of the extent to which the same animals would be selected using these different models in a breeding program where each generation 10% of the animals are selected as parents of the next generation. Also, to understand the relationship between predictions from the models at the genome level, we quantified the overlap between the top 1000 ranked SNP among the models and traits analyzed. For any given trait, an “overlapping SNP” between two models A and B was defined as any SNP in the top 1000 ranked for model A identical or in high LD (r^2^ > 0.90) with a SNP among the top 1000 ranked from model B. This approach may yield different results depending on one starting the comparison from model A to model B or vice versa and, therefore, here we report results for both directions.

### Data and software availability

All data associated with this manuscript can be obtained at https://figshare.com/s/8bdd723be9d0e748cadf. The code developed and used to perform analyzes described in this manuscript are included as Supplementary Material, as well as a detailed description of results. All software used is publicly available.

## RESULTS

### Model performance

The accuracy of predicted phenotypes from GBLUP, BayesB, ENET and GBM for animals in the validation set (generation 11) is shown in Figure 2. The best performing model varied according to the trait being analyzed.

**Figure 2.**
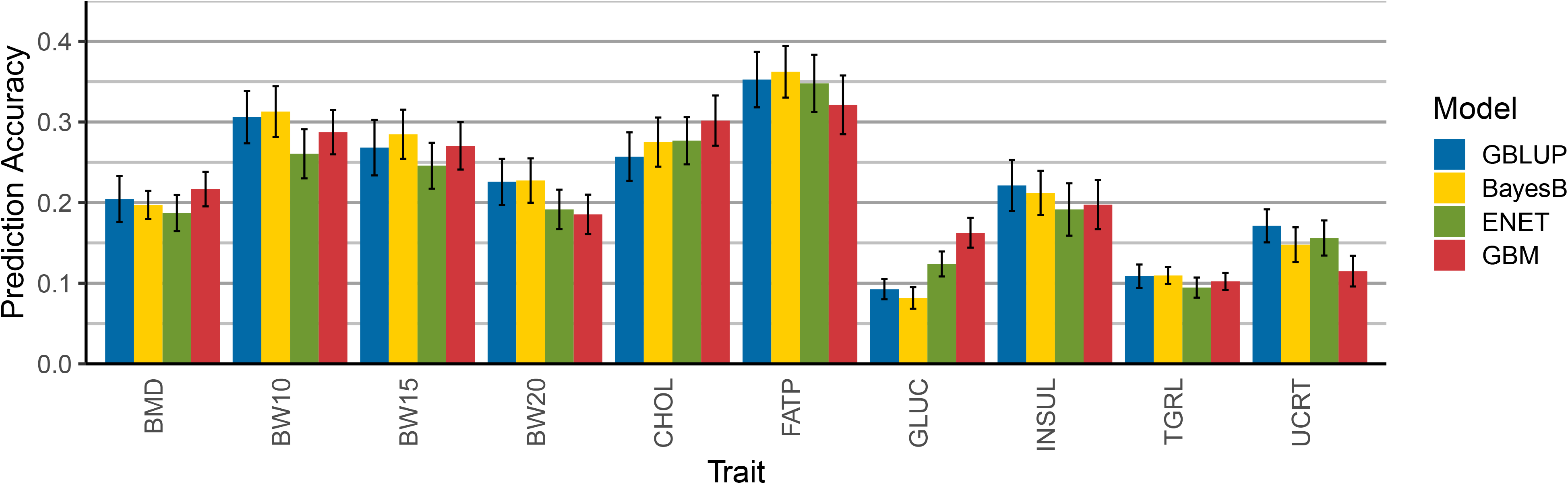
Prediction accuracy, including standard errors, obtained from GBLUP, BayesB, elastic net (ENET) and gradient boosting machine (GBM) for the traits: *bone mineral density at 12 weeks (BMD), Body weight at 10, 15 and 20 weeks (BW10, BW15 and BW20); circulating cholesterol at 19 weeks (CHOL), adjusted body fat percentage at 12 weeks (FATP), circulating glucose at 19 weeks (GLUC), circulating triglycerides at 19 weeks (TRGL), circulating insulin at 8 weeks (INSUL) and urine creatinine at 20 weeks (UCRT)*.

Prediction accuracies obtained for traditional linear models (GBLUP and BayesB) were, in general, proportional to the trait’s heritability, with GBLUP overcoming BayesB for BMD, GLUC, INSUL, TRGL and UCRT. Predictive accuracy obtained with GBLUP was never the worst among tested models for any of the traits. The highest prediction accuracies were observed for body composition traits (BW10, BW15, BW20 and FATP), for which BayesB outperformed all other models. Conversely, BayesB particularly underperformed when analyzing GLUC which was one of the traits with the lowest overall accuracy across linear models. The ENET had lower prediction accuracy when compared to other models across traits. It was never the best performing model for a particular trait and showed the worst performance for BMD, BW10, BW15, BW20, INSUL and TRGL.

The GBM model showed best predictive performance for BMD, CHOL and GLUC. For other traits, prediction accuracy from GBM varied from being competitive to the linear models for BW10, BW15 and TRGL, to a poorer performance observed for UCRT. It only showed the worst predictive ability among all models for FATP, but with a small difference from the next performing model (− 1.76% absolute difference). The GBM model performed particularly well when analyzing GLUC, showing predictive performance much higher than the linear models. Overall, GBM showed a less consistent pattern of predictive performance across trait categories when compared to the linear models.

In terms of prediction error, GBLUP was the model with best performance for most traits, in most cases followed by GBM. The GBM model showed the lowest MSE for BMD, CHOL and GLUC. For all traits, BayesB showed the highest MSE when compared to other models, even for traits for which it had the best prediction accuracy. Relative differences between MSE from the best and worst model were lower for body weight traits (BW10, BW15 and BW20) and higher for CHOL and INSUL.

### Impact of feature selection on prediction performance

Figure 3 shows the prediction accuracy obtained by GBLUP, ENET and GBM when fitting only the top 100, 250, 500, 1000 from a GBM run or all SNPs (52K). When compared to fitting all SNPs (SNPALL), fitting only a subset of important features showed distinct pattern depending on the trait analyzed and model applied.

**Figure 3.**
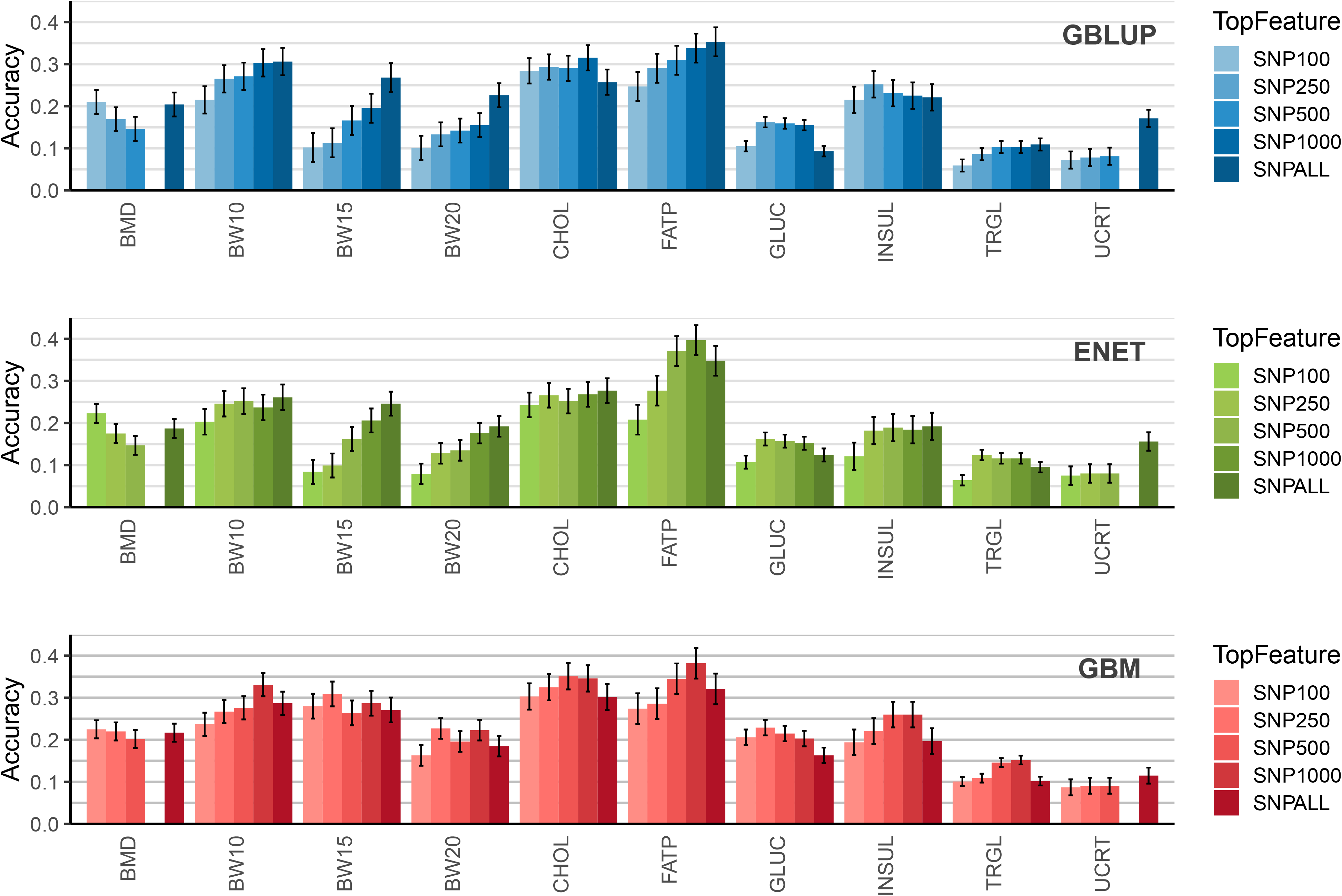
Prediction accuracy, including standard errors, for the analyzed traits for GBLUP (top), ENET (mid) and GBM (bottom) fitting exclusively the top 100 (SNP100), 250 (SNP250), 500 (SNP500), 1000 (SNP1000) ranked by a gradient boosting machine (GBM) model and fitting all SNPs (SNPALL). *Traits: Bone mineral density at 12 weeks (BMD), Body weight at 10, 15 and 20 weeks (BW10, BW15 and BW20); circulating cholesterol at 19 weeks (CHOL), adjusted body fat percentage at 12 weeks (FATP), circulating glucose at 19 weeks (GLU), circulating triglycerides at 19 weeks (TRGL), circulating insulin at 8 weeks (INSUL) and urine creatinine at 20 weeks (UCRT).*

When fitting the GBLUP model, including increasingly more important SNPs resulted, for most traits, in an incremental increase in accuracy, reaching its maximum value in the SNPALL scenario. This was especially the case for traits which were expected to be highly polygenic like BW10, BW15, BW20 and FATP. For CHOL, GLUC and INSUL, fitting GBLUP with a subset of top importance SNPs selected by the GBM model yielded higher accuracy than SNPALL, the number of top SNPs that resulted in the highest prediction accuracy was dependent on the trait being analyzed.

When fitting ENET, including subsets of relevant SNP as predictors for BW10, BW15 and BW20 yielded similar results as for GBLUP. For FATP, there was an incremental increase in accuracy by including more important SNPs, but with SNP500 and SNP1000 showing even higher prediction accuracies than in SNPALL and comparatively higher than the accuracies obtained for FATP by GBLUP. For most other traits (except for BW10 and UCRT), fitting an ENET considering only some top SNPs showed higher prediction accuracies than SNPALL.

The GBM model showed for almost all traits a higher predictive accuracy when considering a subset of SNPs compared to fitting all available SNP (SNPALL). The only exception to that was UCRT, for which the inclusion of important SNPs up to 500 resulted in only a marginal increase in accuracy. For each tested subset of important SNPs, GBM outperformed GBLUP and ENET for prediction accuracy, except for FATP. For this trait, ENET yielded around 0.02 higher absolute accuracy than GBM for SNP1000. For BMD and UCRT, the total number of features selected by GBM was 364 and 419. Consequently, for these traits, running SNP1000 was not possible and SNP500 indicate SNP364 and SNP419, respectively.

### Generation gaps and connectedness between reference and validation sets

Figure 4 shows the prediction accuracies obtained for different scenarios considering increasing distance between reference and validation sets. The increase in distance between the reference and validation sets resulted in a decrease in prediction accuracy for almost all trait/model combinations, in different magnitudes. The exception to that pattern was observed for GLU, for which a marginal increase in accuracy (although not drastically different across scenarios) was observed for GBLUP and GBM. Independent of the trait analyzed or model used, differences in accuracy between NoGAP and GAP9 were much lower than between NoGAP and GAP89 or between GAP9 and GAP89. These differences varied from - 0.20 (BMD – GBM) to +0.03 (GLUC – GBLUP).

**Figure 4.**
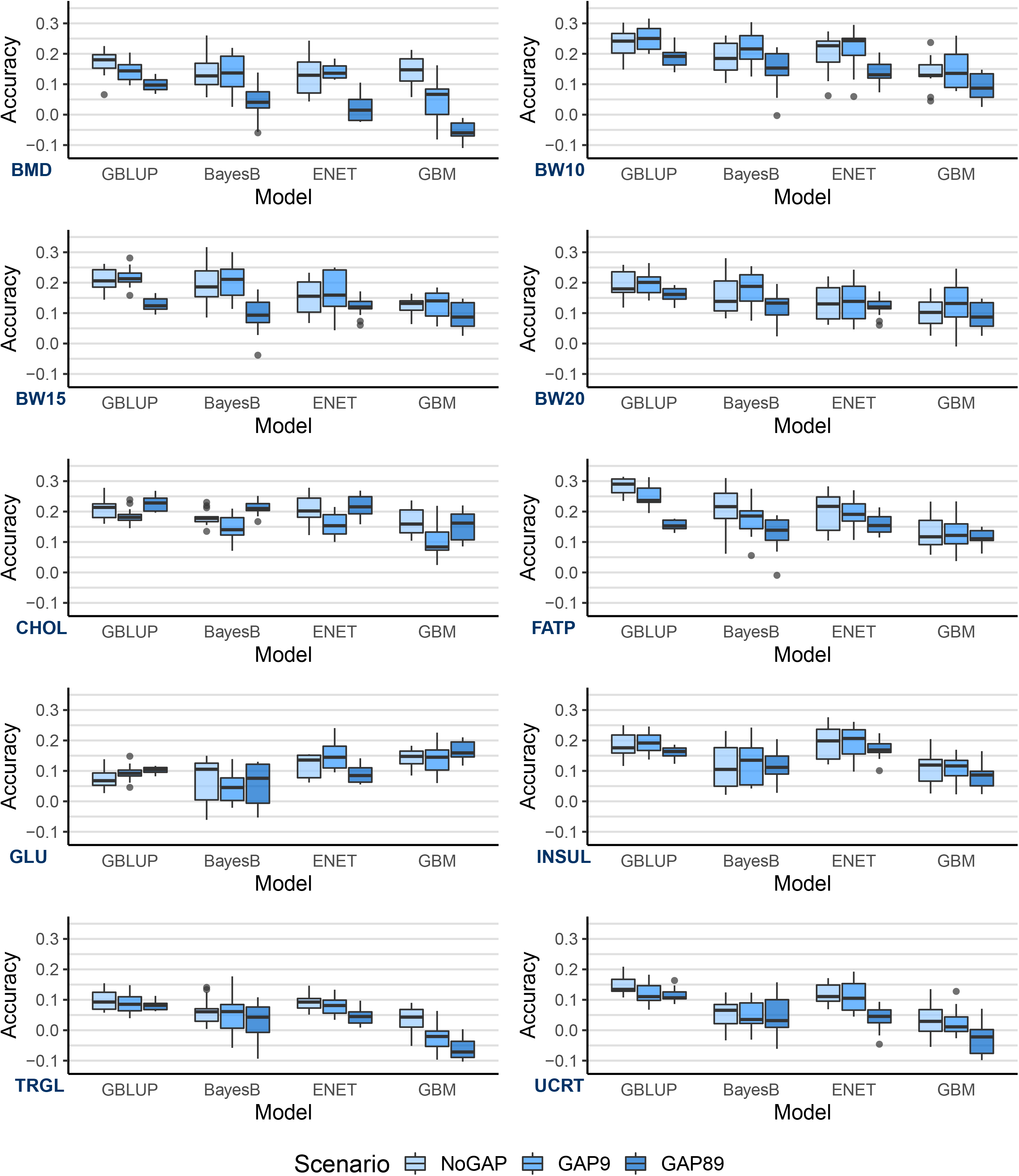
Distribution of prediction accuracies (from 20 replicates) for scenarios including progressive distance between reference and validation sets using GBLUP, BayesB, elastic net (ENET) and gradient boosting machine (GBM) models. *Traits: Bone mineral density at 12 weeks (BMD), Body weight at 10, 15 and 20 weeks (BW10, BW15 and BW20); circulating cholesterol at 19 weeks (CHOL), adjusted body fat percentage at 12 weeks (FATP), circulating glucose at 19 weeks (GLUC), circulating triglycerides at 19 weeks (TRGL), circulating insulin at 8 weeks (INSUL) and urine creatinine at 20 weeks (UCRT)*

The GBLUP model showed the lowest decrease in accuracy between NoGAP and GAP89 scenarios among traits when compared to other models, except for FATP, for which the difference in performance between NoGAP and GAP89 for GBLUP was the highest among all models (−0.12). On the other hand, the GBM model showed the highest drop in accuracy when comparing NoGAP and GAP89 scenario, especially for BMD, TRGL and UCRT. Especially for these traits, using GBM on a GAP89 scenario resulted in negative average prediction accuracies.

Independent of the model used, the traits BW10, BW15, BW20 and FATP showed the lowest decrease in accuracy while BMD, TRGL and UCRT showed the highest decrease in accuracy between NoGAP and GAP89 scenarios. For CHOL the prediction accuracy of GAP89 was higher than observed for GAP9 for all models tested, while for GLU this pattern was observed for predictions from GBLUP, BayesB and GBM, although in smaller differences between scenarios.

The ranking of model accuracy across traits observed using the full dataset (Figure 2) and for the generation gap scenarios (Figure 4) was not the same. When considering the full dataset, GBM yielded the best accuracy for BMD, CHOL and GLU, however the same pattern was not observed for the generation gap scenarios. Overall, when under any of the generation gap scenarios, GBLUP had the best accuracy across traits.

### Animal predictions and SNP ranking similarities between models

The number of unique animals among the top 20 ranked using GBLUP, BayesB ENET and GBM models is shown in Figure 5 (top) for BW10 (A) and GLUC (B). Respectively for these two traits, the number of unique animals in the top 20 rank was 4 and 10 for GBLUP, 10 and 14 for BayesB, 7 and 9 for ENET; and 7 and 11 for GBM. Detailed results for all traits are included in Supplementary Material – Figure S2. Overall, the number of overlapping animals between pairs and triples of models was slightly higher for BW10 than for GLUC. The number of animals uniquely in common between any model and GBM varied between 0 and 4 for BW10 and between 0 and 3 for GLUC.

**Figure 5.**
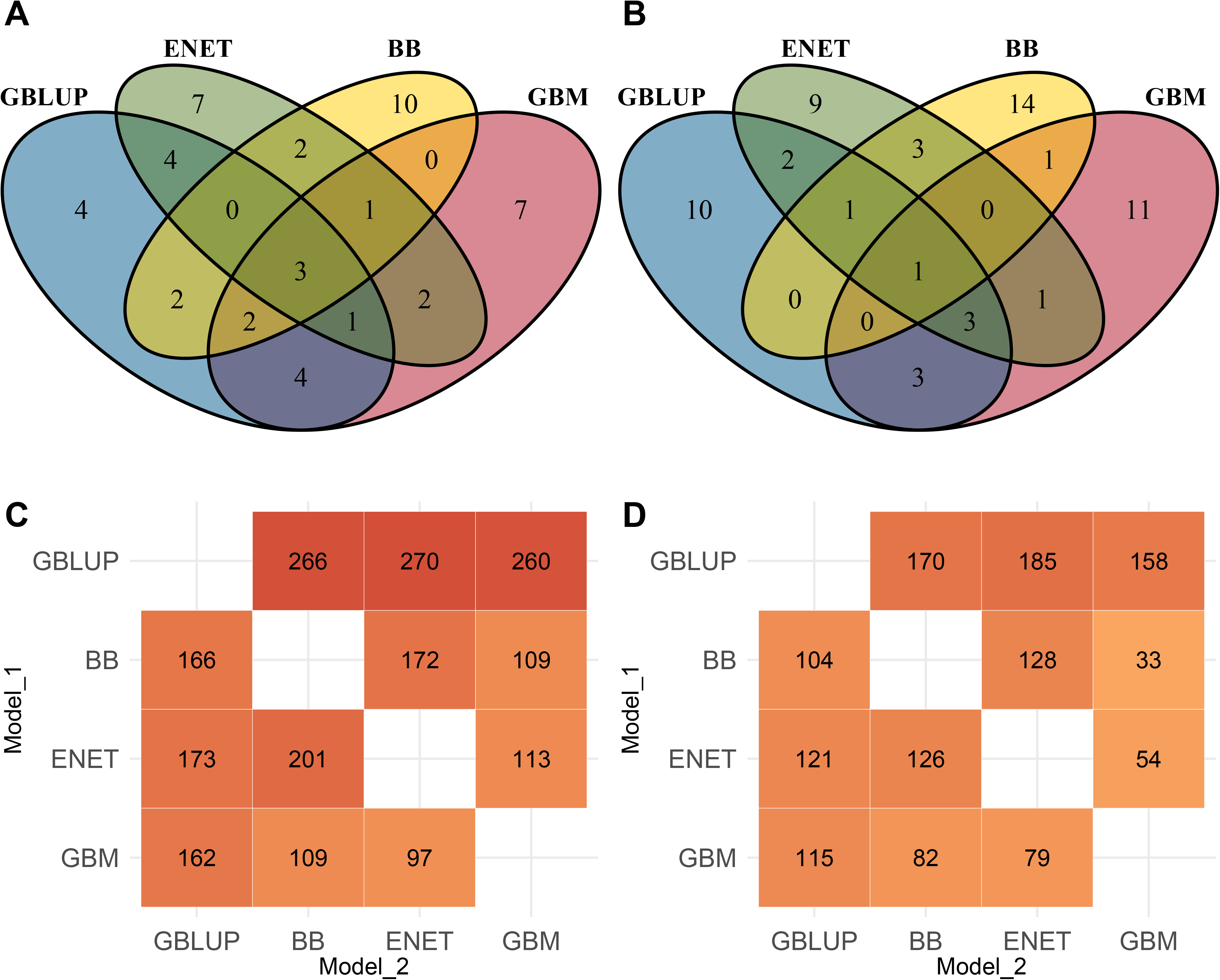
(A and B) Venn diagrams showing the unique animals among the top 20 (above) predicted values (10% of the validation subset) between models and (C and D) the number of SNP markers in common or in high LD (r^2^ > 0.90) among the top 1,000 SNP from GBLUP, BayesB (BB), elastic net (ENET) and gradient boosting machine (GBM) for BW10 (A and C) and GLUC (B and D). In C and D, values represent the overlap of SNP when Model_1 (y-axis) is considered as reference. Traits: *Body weight at 10, weeks (BW10); circulating glucose at 19 weeks (GLUC).*

Figure 5 also shows the count of overlapping markers among the top 1000 ranked by the models investigated for BW10 (C) and GLUC (D). Overall, the number of overlapping markers between any pair of models was higher for BW10 than for GLUC. Within traits, higher values were usually observed for comparisons between two linear models than between a linear model and GBM, while the lowest overlap was observed between ENET and GBM; and between BayesB and GBM. Comparisons between GBLUP and any other model had more overlapping markers than between other models. The largest differences between values above diagonal and the respective comparison below diagonal were observed for comparisons between GBLUP and any other model, with values above the diagonal (GBLUP x other model) being considerably higher than values below the diagonal (other model x GBLUP).

## DISCUSSION

In the present study we compared predictive performances of commonly applied linear methods (GBLUP, BayesB and ENET) and a non-parametric machine learning ensemble method (GBM) for GP of 10 complex phenotypes in the DO mouse population. Although the evaluation of feasibility of genomic selection in mice was not our focus, results of predictive accuracy can be used as a guide if selection is intended for this population. Currently, the mating scheme used for the DO population is a randomized outbreeding strategy (Churchill et al., 2012), however, being able to predict phenotypes could be useful if any directional selection is of interest in the future.

Accuracies of GP have been reported by previous authors in another mice population (Legarra et al., 2008; Lee et al., 2008). Overall results showed low to medium predictive accuracies, ranging from 0.10 to 0.65 depending on the trait analyzed and cross-validation strategy considered. Our results confirmed that the performance of genomic prediction methods seem to be highly dependent on the trait’s genetic architecture. When analyzing the traits that are mostly polygenic (BW10, BW15, BW20, FATP and TRGL), linear models were able to outperform GBM in both the full dataset (Figure 1) and for scenarios with lower connectedness between reference and validation subsets (Figure 4). BayesB was the best model for the three BW traits and FATP, while GBLUP had the best results for TRGL. In a genome-wide study using data from the same population, Zhang et al. (2010) showed an absence of QTL with pronounced effects for TRGL, with mostly small effects detected for genome-wide markers. This could explain why GBLUP had better predictive performance than BayesB or ENET for this trait.

Among the ten traits analyzed, evidence of non-additive effects has been reported for BMD (Tyller et al. (2016), CHL (Stewart et al., 2010; Li and Churchill, 2010) and GLU (Stewart et al., 2010; Chen et al., 2017). Coincidently for these traits GBM showed a better predictive performance than the linear models in the full dataset. Based on their results in strawberry using convolution neural networks, Zingaretti et al. (2020) suggested that machine learning methods may outperform parametric and semi-parametric models when the epistatic component is relevant (proportionally to the additive genetic variance) and narrow-sense heritability is medium to low (below 0.35). This is roughly in line with our results for CHL (h^2^ = 0.33), GLU (h^2^ = 0.11) and BMD (h^2^ = 0.39). Interestingly, in our results the superiority of predictive ability from GBM compared to the parametric models was higher for the trait with lower heritability (GLU) than for CHL and BMD. Low-heritability traits imply that a smaller portion of observed variance is explained by the additive component, and therefore, any other non-linear effects might explain proportionally more of the phenotypic variance than in high-heritability traits. This larger proportion of the phenotypic variance with a non-linear origin can more easily be captured by the GBM model, increasing performance of the model for such traits. Overall, the observed ranking of model performance across anticipated trait architecture was in line with previously reported results. In a detailed simulation study, Abdolahi-Arpanahi et al. (2020) showed that for traits controlled by many QTL (1000) with only additive effects, GBLUP and BayesB outperformed any machine learning approach, while for traits controlled by a small number of QTL (100) with non-additive effects, GBM largely outperformed other parametric and non-parametric models. Note that in their study, traits were simulated with only additive or non-additive effects, which is not expected to be the case in real world situations. However, their results on these extreme cases, are a robust indication of what to expect from each type of genomic prediction model. The similarity between results obtained in the present- and the afore-mentioned studies are in line with the current knowledge of genetic architecture of the analyzed traits (Table 1).

The efficient built-in feature extraction from GBM enables pre-screening of SNPs (Lubke et al., 2013; Li et al., 2018); and, therefore, minimize the loss in accuracy when reducing the number of markers in a genotype panel. The performance of GBM on pre-selection of informative SNP markers varied across traits and models subsequently used for phenotype prediction. When considering the highly polygenic traits (BW10, BW15, BW20, FATP and TRGL), using pre-selected SNP markers generally decreased accuracy of GBLUP. However, for ENET and GBM, in certain situations a subset of pre-selected SNP tended to yield higher predictive accuracy than using the complete SNP panel (Figure 3). For traits with evidence of non-linear effects (BMD, CHL and GLU), a similar pattern was observed, with the difference that the use of subsets of markers more commonly resulted in higher predictive accuracy than when fitting the models with all available SNP. After pre-selection of informative markers, GBM showed the biggest gains in accuracy across traits and models, which is expected, since we used a GBM model to accomplish the former. Azodi et al. (2019) observed that feature selection (using the random forest method) notably improved prediction accuracies when using artificial neural networks (ANN) in multiple plant species. However, in their case, predictive accuracies using ANN were overall lower than other models. Using data from Brahman cattle, Li et al. (2018) investigated the potential of three different ensemble learning methods to pre-select SNPs and showed that GBLUP accuracies using SNPs preselected with GBM in some cases were actually similar to accuracies based on all SNPs. Together with our findings, the above-mentioned results suggest that GBM can be used for pre-screening informative markers, even when further genomic prediction is performed using traditional linear models, such as GBLUP. One limitation of ours and all investigations found in literature is the focus in performing feature selection and further fitting top relevant markers into univariate models. Further research is needed to expand this from a univariate to multivariate approach for practical implementation in genomic selection breeding programs.

Curiously, for UCRT the inclusion of pre-selected SNP (from 100 to 500) did not affect predictive accuracy, which was similar across scenarios and models, but always lower than using the full SNP panel. This may occur because the optimum number of informative markers might be above 500 or just that GBM was not successful at pre-selecting informative markers for this particular trait. A similar pattern was previously reported by Azodi et al. (2019) when fitting different numbers of informative pre-selected markers into a model for genomic prediction in sorghum. Authors observed low and stable prediction accuracy (around 0.40) when using up to 5% of top markers, but a strong increase when using more than 5% of top relevant markers, reaching up to 0.60 when using 80% of available markers. We have replicated the feature selection of top 100, 250, 500 and 1000 SNPs using BayesB instead of GBM. Results suggest a superiority of GBM for pre-selecting informative markers (Supplementary Material – Figure S1) as predictive accuracy across traits was consistently lower when using BayesB compared to using GBM for the same task.

The size of the reference population and the strength of the connectedness between reference and validation subsets have been shown to influence GP accuracies from linear models (Habier et al., 2007; Wientjes et al., 2013; Liu et al., 2015). In terms of connectedness, maximizing predictive performance involves maximizing connectedness between reference and validation populations, while simultaneously minimizing connectedness within the reference population (Pszczola et al., 2012). Although extensive research has been done over this topic regarding traditional GP using parametric models, this is not the case for ML models. In addition to that, much has been discussed in literature about how “data-hungry” machine learning models could be. However, studies have not only shown no clear superiority of predictive performance from machine learning over parametric models when using large datasets (Bellot et al., 2018), but also good performance of the same machine learning models when using datasets of hundreds of individuals (Azodi et al., 2019; Zingaretti et al., 2020; Bargelloni et al., 2021). When compared to the predictive performance of linear models, GBM had competitive results for most traits and a superior performance for BMD, CHL and GLU when using the full dataset (Figure 2). However, this relatively good performance was not maintained for NoGAP, GAP9 and GAP89 scenarios that contained less data (Figure 4). This pattern was observed across all traits and scenarios and may indicate that using only 300 individuals in the reference subset affected more drastically the predictive performance of the GBM model than GBLUP, BayesB or ENET. Overall, the decrease in accuracy observed from NoGAP to GAP89 was also more severe for GBM than for other models. We hypothesize that this could happen because as the distance between reference and validation populations increases, the frequency of recombination events also increases between genotypes from individuals in the two subsets. As GBM implicitly fits SNPxSNP interactions, the increased number of recombinations will impair the accurate estimation of allele combinations and interactions.

The ultimate aim of genomic prediction in the breeding context is to make accurate selection decisions early in the animal’s life. Therefore, comparing the top ranked individuals between methods is a useful way to understand how different these are in practical terms. In the present study, independent of the trait analyzed, linear models shared many more individuals among the top 20 best from the three models (GBLUP, BayesB and ENET) than with GBM. For GLUC, for which we expected non-additive effects, the similarity between rankings for linear models was lower, while the number of unique animals for a single model were higher. On the other hand, as we consider BW10 to be controlled mostly by additive effects, the absence of relevant non-additive effects is probably the cause of lesser differences between linear models and GBM regarding selection decisions.

We evaluated the overlap among top ranked SNP between the different models (Figure 5, Supplementary Material – Figure S3). One thing that must be acknowledged is that there are differences in the way each of the different models estimate the relevance of a single SNP. This may affect the comparison of the overlapping relevant genomic regions between methods for a certain trait. For the linear models, SNP relevance is based on changes observed at the phenotypic level by the change in allelic dosage (0,1,2), while for GBM a SNP is considered relevant when the inclusion of this SNP in the decision tree contributes to a reduction in prediction error, and this can be affected by other SNP also used in the same decision tree. On the other hand, when used for genomic prediction, these differences will impact the obtained genomic predictions and thereby indirectly impact selection decisions. Therefore, this simple comparison of SNP ranks is informative to understand the similarity of outcomes from different models.

The asymmetry of results obtained from the overlapping top ranked SNP between models can be seen comparing values below and above diagonals in Figure 5 (C and D). The strongest driver of the differences observed seems to be the ability of models to perform variable selection. When starting comparisons from GBLUP (first row above diagonals in Figure 5 - C and D), there were many SNP located in specific short genomic regions among the top 1000 ranked SNP for this model. Several top markers from GBLUP were in high LD with at least one top ranked marker from the other models. In contrast, the variable selection applied by BayesB, ENET and GBM, resulted in fewer SNPs within a given genomic region to be among the top ranked ones. As a consequence, the number of top ranked SNP in high LD with top ranked SNPs from the other models was much lower. Therefore, the difference between values above and below diagonal are directly related to the difference in magnitude of penalization applied to markers between any given pair of models. When comparing results from genomic prediction of height in maize using BayesA, ENET and random forest models, Azodi et al. (2019) have observed marked dissimilarity among the top 8000 markers. Results showed that BayesA and ENET shared 1589 (20%) markers, while RF shared 328 (4%) markers with BayesA and 475 (6%) with ENET. In the present study, this higher similarity among SNP ranks between linear models in addition to much lower similarity between linear models and an ensemble machine learning model (random forest in Azodi et al. [2019] or GBM in the present study) was also observed for BW10. At the same time, the difference between average SNP overlaps between two linear models or between a linear model and GBM was much lower for GLUC. From these results we can hypothesize that linear models have similar SNP rankings for polygenic traits because the underlying genetic architecture is in line with assumptions and parametrization considered in such models, while the presence of non-linear effects is probably captured differently by the distinct linear models, generating the observed overall dissimilarity.

## CONCLUSION

Gradient boosting machine had a competitive performance for genomic prediction of complex phenotypes in mouse specifically for traits with non-additive effects where it can outperform linear models. The gradient boosting machine was more affected by datasets with less data points and by decrease in relationship between reference and validation populations than linear models. Considerable differences between the top ranked animals suggest that using linear models versus GBM will result in clear differences in selection decisions. The built-in feature selection from GBM seems beneficial to extract a smaller number of informative markers and in some cases can improve accuracies even when parametric models are used for prediction.

